# Large-scale calcium imaging with a head-mounted axial scanning 3D fluorescence microscope

**DOI:** 10.1101/2021.01.20.427512

**Authors:** Yuichiro Hayashi, Ko Kobayakawa, Reiko Kobayakawa

## Abstract

Miniaturized fluorescence microscopes are becoming more important for deciphering the neural codes underlying various brain functions. With gradient index (GRIN) lenses, these devices enable recording neuronal activity in deep brain structures. However, to minimize any damage to brain tissues and local circuits, the diameter of the GRIN lens should be 0.5–1 mm, resulting in a small field of view. Volumetric imaging capability might increase the number of neurons imaged through the lenses considering the three-dimensional (3D) structure of neural circuits in the brain. To observe 3D calcium dynamics, we developed a miniaturized microscope with an electrically tunable lens and a novel CNMF-based neural signal extraction algorithm for wide-field 3D imaging data. By combining the hardware and software, approximately 1000 neurons were imaged from the cortices of freely behaving mice. Compared with the state-of-the-art 2D imaging technique, the proposed 3D method imaged 1.7–2.6 times more cells with a higher separation of cellular signals.

## Introduction

Calcium imaging enables to record the activity of hundreds of neurons at cellular resolution simultaneously. In particular, the combined use of miniature head-mounted microscopes and genetically encoded calcium indicators allows tracking the neuronal activity for days to months in freely moving small animals ^1,2^. For deep brain imaging, gradient index (GRIN) lenses have been used for deep brain imaging to relay images to microscopes ^1–4^. However, the implantation of GRIN lenses damages brain tissue. Lenses with a smaller diameter (0.5–1 mm) and a small field of view might be used for deep brain imaging to minimize the damage. Considering the three-dimensional (3D) structure of neural circuits in the brain, 3D imaging represents a possible strategy to maximize the number of recorded cells through a limited field of view.

Light-field microscopy (LFM) has performed 3D fluorescence calcium imaging ^5,6^. Typically, LFM maps a 3D image onto an image sensor using a microlens array. The 3D image is then computationally reconstructed from the 2D image captured by the image sensor ^7^. A miniature implementation of LFM, MiniLFM, imaged fluorescence calcium signals within a volume of 700 × 600 × 360 μm^3^ at near single-cell resolution in the hippocampal CA1 area of freely moving mice ^8^. In LFM, spatial and depth data are mapped onto a single image sensor. Therefore, the LFM frame rate can be as high as 2D imaging, but the spatial resolution is usually low. To overcome this limitation, Miniscope3D employs an optimized microlens array and a new reconstruction algorithm ^9^. The system achieves ∼2× higher spatial resolution than that of MiniLFM. However, its calcium imaging performance *in vivo* calcium has not been evaluated.

An alternative strategy for 3D imaging is to add a focusing element to the microscope and sequentially capture 2D images at different depths. The electrowetting lens (EWTL) is a small, fast, and lightweight variable focus lens. EWTL-based 3D imaging systems combined with a one- or two-photon laser scanning microscope have been used in small animals ^10,11^. In these systems, the head-mounted optical unit contains only the objective lens. The EWTL connects to the laser scanning device with an optical fiber bundle. Therefore, the head-mounted unit is lightweight. However, the optical fiber bundle limits the spatial resolution of the system. Moreover, these systems rely on point scanning, which restricts the scan rate of 3D volumes.

Owing to the high and fluctuating background, extracting neuronal signals from wide-field imaging data is another challenging issue in 2D and 3D imaging. Principal component analysis/independent analysis (PCA/ICA) was the first attempt to extract neuronal signals from wide-field imaging data automatically ^12^ and has been applied to 3D imaging ^6^. However, it is a linear demixing method and therefore often fails to decorrelate signals from spatially overlapping cells ^13^.

Sparse matrix factorization ^14^ and non-negative matrix factorization (NMF) ^15^ overcome this problem. These methods can deal with overlapping cells by introducing nonlinearity. Constrained NMF (CNMF) improved the demixing performance by introducing temporal dynamics of the calcium activity ^13^. However, the original CNMF implementation was designed for images with low-background fluorescence, such as two-photon and light-sheet microscopy. The algorithm performance is often poor on wide-field imaging data ^16^. Although several CNMF-based methods have been proposed for handling wide-field imaging data ^17,18^, they are unsuitable for 3D imaging.

Here, we developed an EWTL-based wide-field 3D fluorescence microscope and a novel CNMF-based neuronal signal extraction algorithm termed CNMFw3 to record the calcium activity from freely behaving small animals at cellular resolution. Combining the hardware and software, we imaged ∼1000 neurons from the cortex of freely behaving mice with a higher separation of cellular signals than that obtained with conventional 2D imaging.

## Results

### Microscope design

We designed a miniature microscope to capture 3D images from freely behaving small animals. Because of its accessibility and low cost, the design was based on the UCLA miniscope v3 (miniscope.org). An EWTL was inserted in the optical path between the objective lens and the dichroic mirror for axial scanning (Figure 1A). The microscope body was 3D-printed to allow rapid prototyping, and the parts were reusable ^19^. The weight of the microscope was 3.7 g, allowing the mice to run freely (Figure 1B). We used several types of objective lenses for different applications. Indeed, the aspherical singlet (Asph; Figure 1C, *Left*) is suitable for relay optics because of its short length and high numerical aperture. A long focal length lens was used for large fields of view (Ach; Figure 1C, *Center*). We used tandem achromatic doublets (TAch; Figure 1C, *Right*) to image the tissue through a cranial window because of their small tip diameter (2 mm), which allows access to the small cranial window on the tissue surface (Figure 1G). The axial scan range with these lenses was 300–600 μm (Figure 1D). Figure 1E and Supplementaly Table 1 show the optical performances. The microscope captured volumetric images at 5 Hz, and each volume image consisted of 12 slices (Figure 1F). The last image in the Z-scan sequence was blurry because of the limited scanning speed of the EWTL (Figure 1G). Therefore, 11 of 12 slices were used to reconstruct the 3D image.

**Figure 1.**
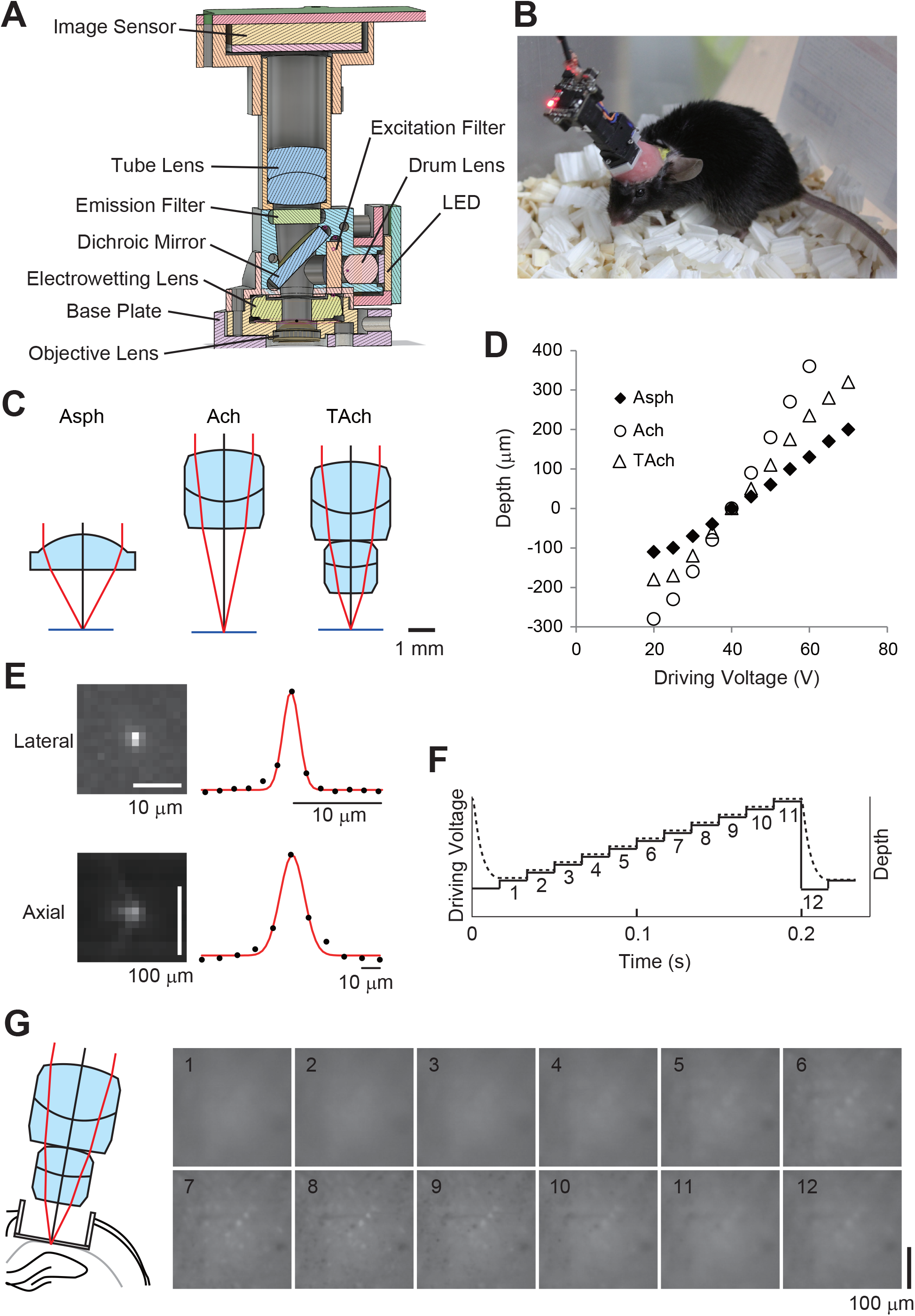
3D fluorescence microscope. **(A)** Section diagram of a 3D microscope. **(B)** Photo of a mouse with the 3D microscope. **(C)** Representation of the three types of objective lenses. *Left*, aspherical singlet (Asph). Center, achromatic doublet (Ach). *Right*, two achromatic doubles in tandem (TAch). **(D)** Relationship between the driving voltage and the depth of focus. **(E)** Optical resolution of the microscope equipped with the aspherical singlet lens. Images and profiles of the 1-μm fluorescent beads. Red lines indicate Gaussian fit. **(F)** Image acquisition sequence. The solid line indicates the driving signal for the ETWL. The dashed line indicates the actual depth of focus. **(G)** Image examples for each Z-slice. The images were taken from the mouse hippocampal CA1 area using the tandem achromatic doublet lens.

### CNMF-based neuronal activity extraction algorithm for wide-field 3D imaging data

To extract images from the wide-field 3D microscope, we developed a new algorithm based on CNMF. Our new method, termed CNMFw3 (CNMF for wide-field 3D data), first estimated the background of the raw image stream with rank-1 matrix factorization (Figure 2A,B). The background-subtracted images were then spatially high-pass-filtered to remove the remaining background (Figure 2C). Next, we looked for initial cell candidates employing the greedy initialization method used in CNMF-E ^18^ with modifications to fit 3D data. The method seeks the local maxima of the product of local correlation image and peak-to-noise ratio image (Figure 2D, E). Finally, the initial spatial and temporal components were refined with CNMF ^13^ (Figure 2F, G).

**Figure 2.**
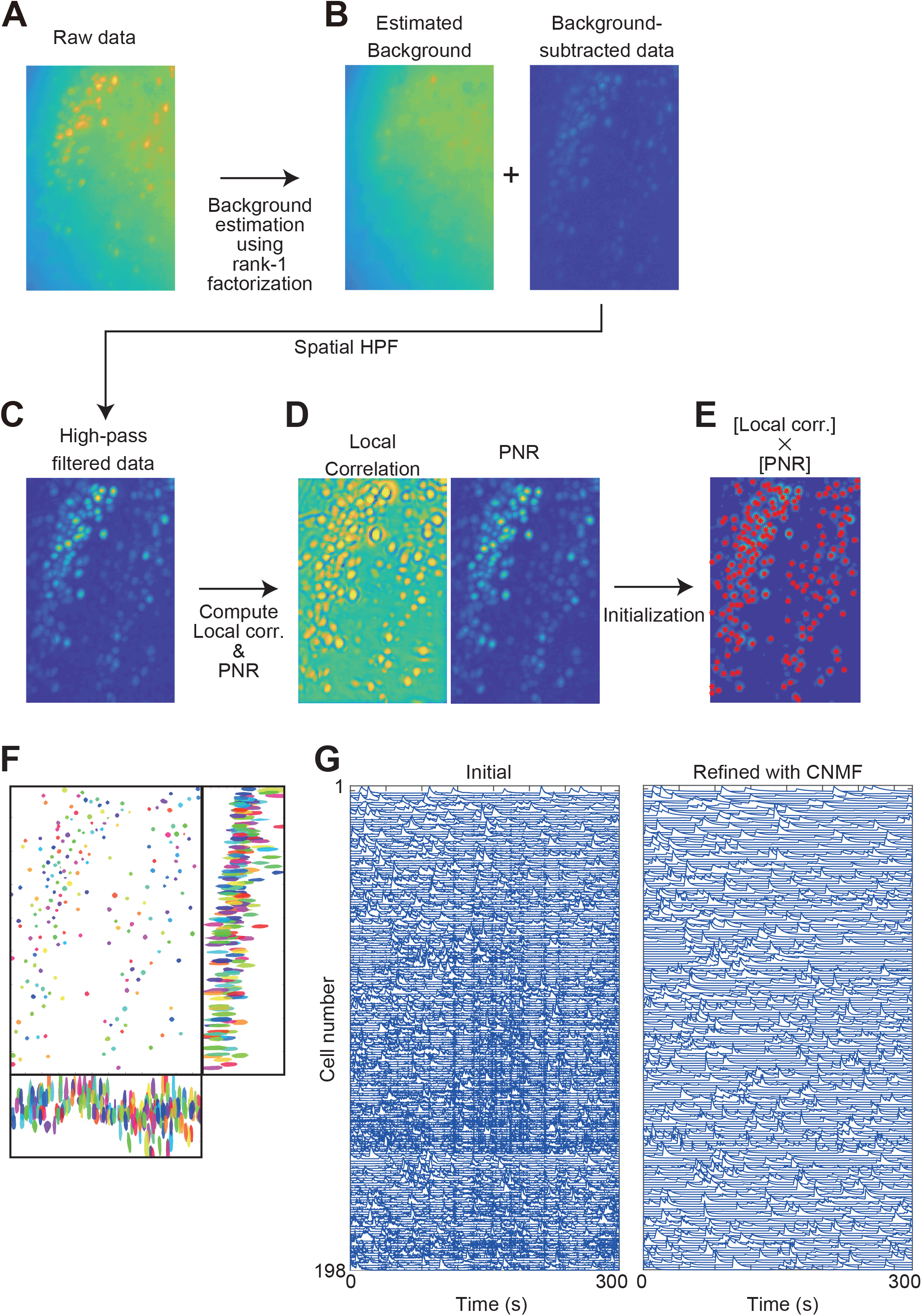
CNMFw3 algorithm. **(A)** Raw 3D image. Images in panels A–E are maximum intensity projections of z- and t-stacks. **(B)** Background estimation by rank-1 factorization. **(C)** Spatial high-pass filtering. **(D)** *Left*, map of temporal correlation between the filtered temporal traces of each pixel and those of neighboring pixels. *Right*, the peak-to-noise ratio of the filtered traces of each pixel. **(E)** Local maxima of the product of local correlation image and peak-to-noise ratio image. Red asterisks indicate the local maxima as the center of initial cell candidates. **(F)** Contours of all neurons were detected by CNMFw3 on three orthogonal planes. The contours of 80% of the peak values of each neuron are displayed. **(G)** Fluorescence traces of initial cell candidates (*Left*) and cell candidates refined with CNMF (*Right*).

### CNMFw3 accurately recovers neuronal activity in simulated data

We compared the neuronal activity extraction performance of CNMFw3 with that of other algorithms for 3D imaging data, namely PCA/ICA and CNMF (plain CNMF) and CNMF-E, the most common ones widely used algorithm for 2D wide-field imaging data. We synthesized 4 image streams with various signal-to-background ratios (SBR, defined as the amplitude ratio between the signal and the background) ranging from 0.1 to 0.8. Each image stream (100 × 100 × 10 pixels, 1000 frames) contained 100 synthesized neurons and background fluctuations extracted from real imaging data (Figure 3A). Poisson noise was added to each pixel to simulate photon shot noise. The number of well-extracted neurons (temporal correlation to the ground truth > 0.9) was higher using CNMFw3 than that obtained with the other methods (Figure 3B,C). PCA/ICA and plain CNMF showed poor performances at the low-SBR range (SBR = 0.4). The performance of CNMF-E was excellent even at SBR = 0.4. However, the number of well-extracted neurons was approximately half of that acquired with CNMFw3 because CNMF-E only handles one Z-slice of the entire volume. Notably, unlike other methods, CNMFw3 still extracted cells at SBR = 0.2 (Figure 3B), indicating that the method can extract more neurons than other methods in real data.

**Figure 3.**
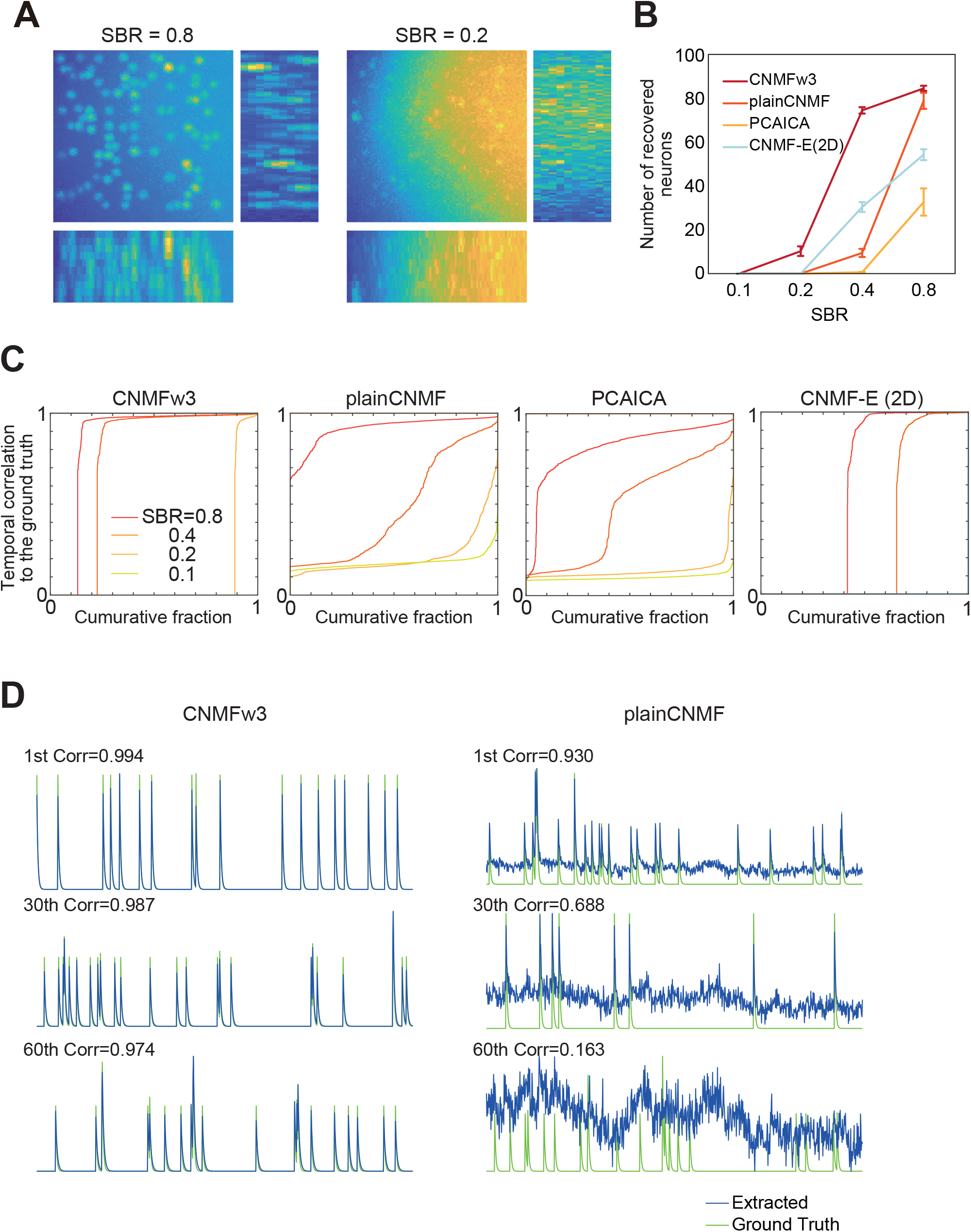
Performances of CNMFw3. **(A)** Representative images of simulated data for high (0.8) and low (0.2) SBR. Images are the maximum projection of an image volume on three orthogonal planes. **(B)** Number of recovered cells from the simulated data using four methods. The number of cells with a high correlation between their fluorescence trace and the ground truth (>0.9) are plotted. The data represent mean ± SEM (n = 10). **(C)** Identification accuracy of the four methods. Correlation coefficients of the recovered trace to the ground truth are shown. **(D)** Recovered traces for intermediate SBR (0.4) dataset (blue) and the corresponding ground truth traces (green) are shown. The 1st, 30th, and 60th from the highest correlation to the ground truth are displayed.

### Application of CNMFw3 to *in vivo* imaging in freely behaving mice

We next applied the hardware and software to *in vivo* microendoscopic imaging. We imaged neurons in the mouse prefrontal cortex (PFC) through 1.0 mm diameter GRIN lens (Figure 4A, B). Volumetric fluorescence images of GCaMP6f-expressing neurons in the PFC were processed with CNMFw3 (Figure 4C, D). CNMFw3 extracted 973 putative neurons. Figure 4E shows fluorescence traces extracted from the 3D movie.

**Figure 4.**
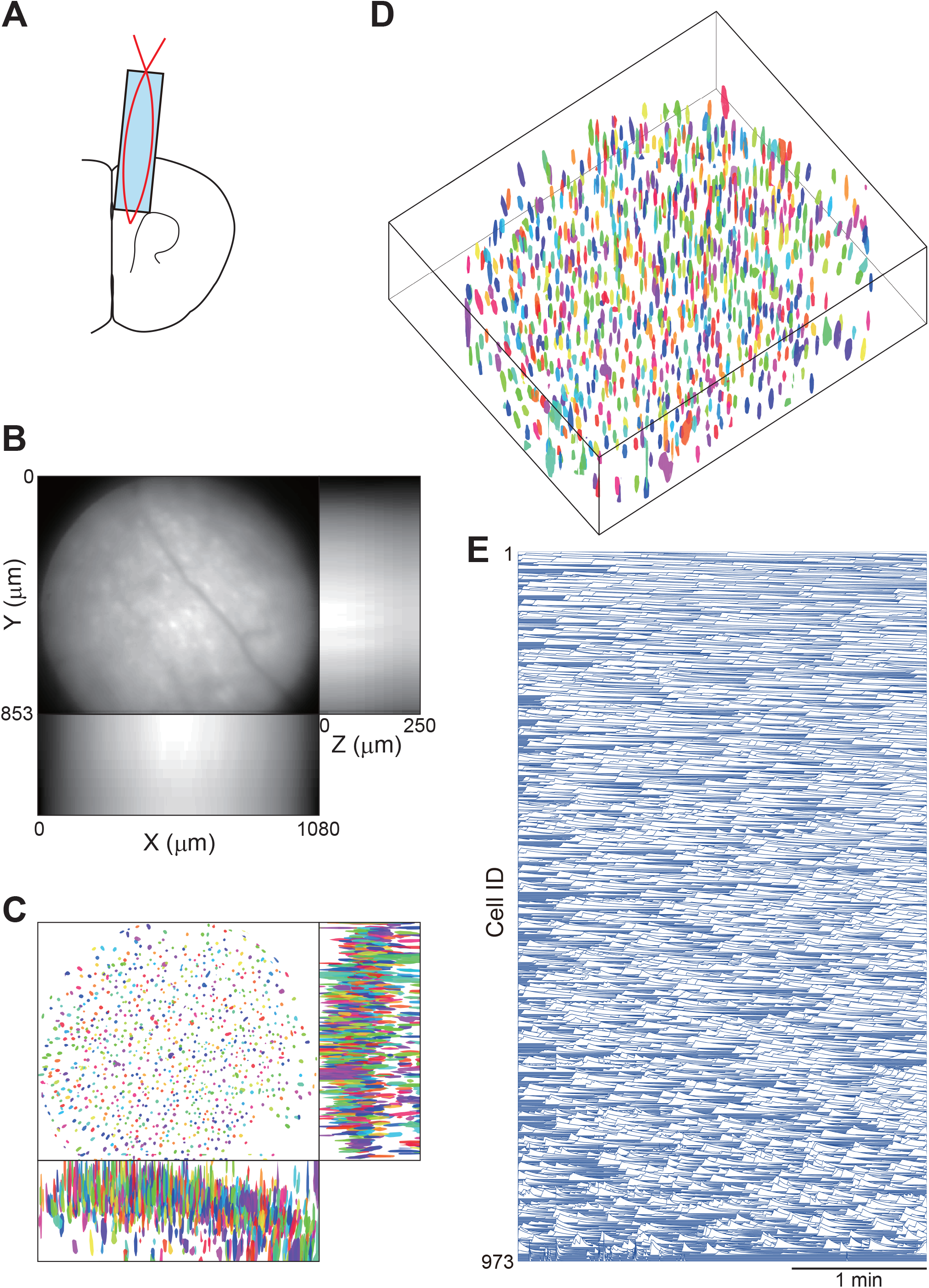
Three-dimensional fluorescence calcium imaging of PFC neurons in a freely behaving mouse. **(A)** Scheme of the anatomical location of the optical path. A GRIN lens (1.0-mm diameter, 3.7-mm length) was inserted in the PFC. **(B)** Maximum projection of an image volume on three orthogonal planes. **(C)** Contours of all neurons detected by CNMFw3. The contours of 80% of the peak values of each neuron are displayed. **(D)** Isometric perspective of data shown in **(C)**. **(E)** Fluorescence traces extracted by CNMFw3.

3D imaging is more beneficial for deep brain imaging because the smaller diameter GRIN lens minimizes tissue damage when imaging deeper brain structures. Therefore, we used the 3D microscope to image the mouse central amygdala (CeA), located approximately 4 mm below the brain surface, through a 0.5-mm diameter GRIN lens (Figure 5A, B). In total, 130 neurons situated at different depths were visible, as shown in Figure 5C and D, and their fluorescence traces were extracted by CNMFw3 (Figure 5E).

**Figure 5.**
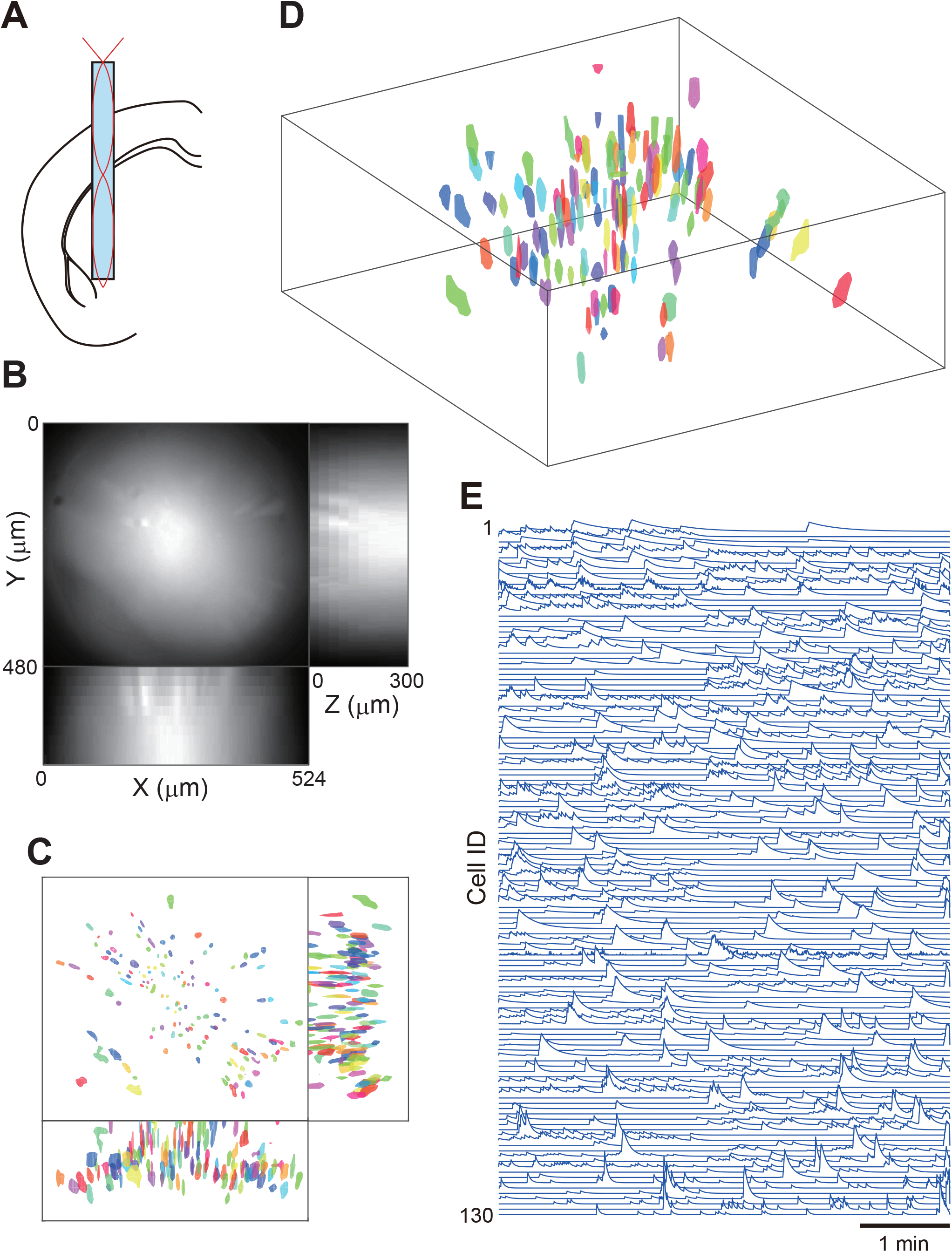
3D calcium imaging in the CeA. **(A)** Scheme of the anatomical location of the optical path. A GRIN lens (0.5-mm diameter, 7.4-mm length) was inserted in the CeA. **(B)** Maximum projection of an image volume on three orthogonal planes. **(C)** Contours of all neurons detected by CNMFw3. The contours of 80% of the peak values of each neuron are displayed. **(D)** The isometric perspective of data shown in (C). **(E)** Fluorescence traces extracted by CNMFw3.

### Advantages of 3D over 2D imaging

To evaluate the merit of the proposed 3D imaging technique in real data analysis, we compared it with conventional 2D imaging. We used CNMF-E, a popular algorithm for wide-field 2D imaging data ^18^, to extract neural activity from 2D images. The number of cells extracted from 2D imaging was defined as the largest number of cells extracted from single Z-slices in the 3D volume (Figure 6A). Significantly more cells were detected from the 3D image than from the 2D image in the PFC and CeA (Figure 6B,C; *t* = 19.54, *p* = 0.0026, n = 3 mice for the PFC, *t* = 3.278, *p* = 0.031, n = 5 mice for the CeA; one-sample two-tailed *t*-test).

**Figure 6.**
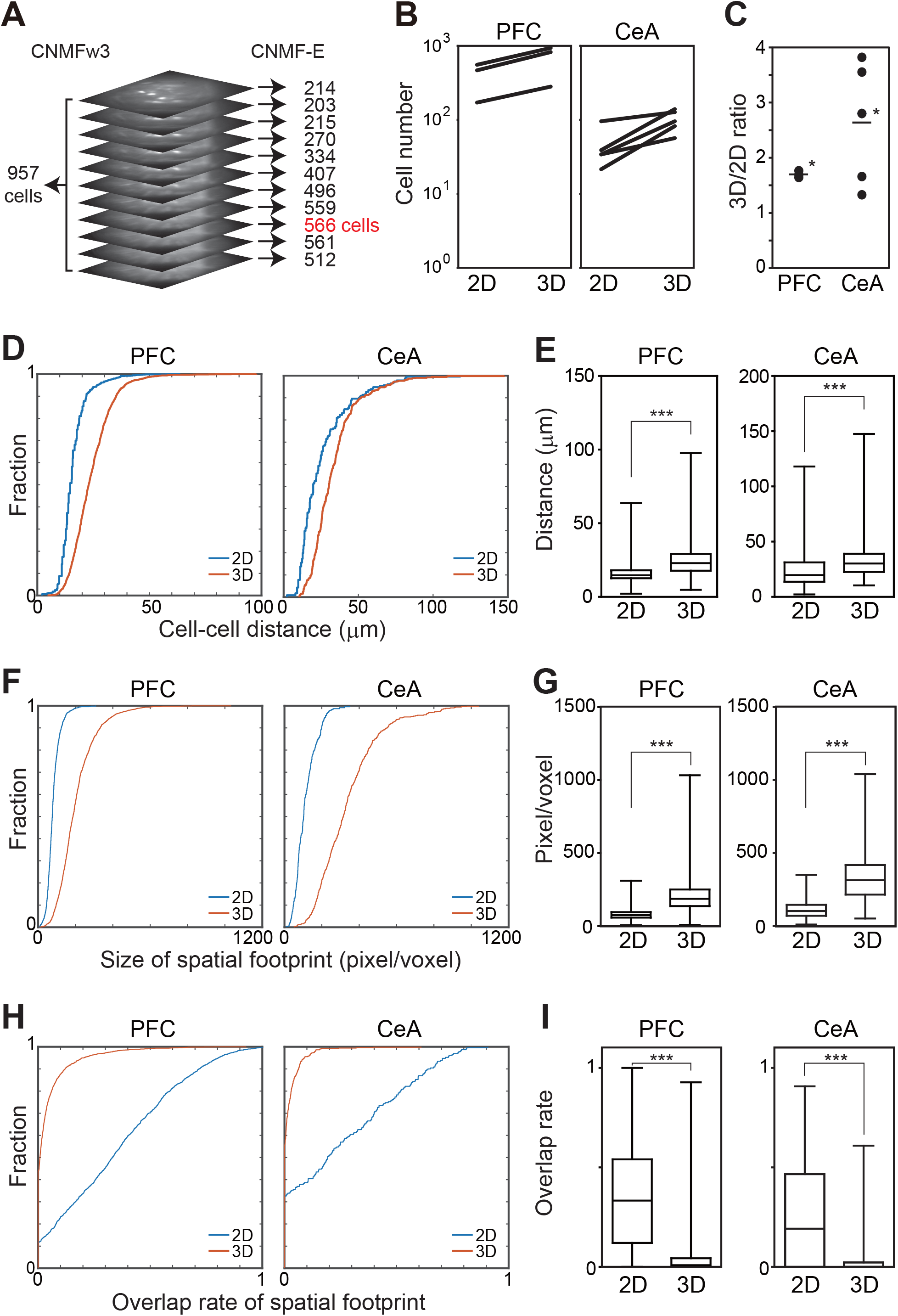
Comparison between 3D and 2D imaging. **(A)** Experimental design. Recorded 3D images were processed with CNMFw3 (*Left*), and each Z-slices in the 3D image were processed with CNMF-E (*Right*). The number of cells extracted from the 2D image is defined as the largest number of cells extracted from a single Z-slice (shown in red) in the 3D volume. **(B)** Comparison of the number of cells extracted from the 3D and 2D images. **(C)** The ratio between the cell numbers extracted from the 3D image and that from the 2D image. The data represent the individual (solid circles) and mean (bars) values. **(D)** Cumulative distribution of the minimum cell-cell distances in 2D and 3D. The cell-cell distance was defined as the distance between the center of mass of each spatial footprint. **(E)** Box-whisker plot indicating the median, quartiles, and range of minimum cell-cell distances. **(F)** Cumulative distribution of the size of the spatial footprints in 2D and 3D. **(G)** Box-whisker plot indicating the median, quartiles, and range of the size of spatial footprints. **(H)** Cumulative distribution of the overlap rate of spatial footprints in 2D and 3D. **(I)** Box-whisker plot indicating the median, quartiles, and range of the overlap rate of spatial footprints. Asterisks indicate significant differences. \**p* < 0.05, \*\*\**p* < 0.0001.

In wide-field 2D imaging, neurons in, above, and below the focal plane are recorded. This is problematic for extracting single-cell activity from the image because signals from multiple neurons tend to overlap. Although recent methods for neural activity extraction partially allow demixing such overlapping signals, reducing the overlap will improve the accuracy of the neural activity extraction ^13,18^. To test whether our 3D imaging method obtained a reduced cell-cell overlap, we compared the minimum cell-cell distance and overlap of the spatial footprint of extracted cells. As shown in Figure 6D and E, the minimum distance between neighboring cells was significantly larger with 3D imaging than that in 2D (*U* = 489100, *p* < 0.0001, n = 2083 cells for 3D imaging of the PFC, n = 1227 cells for 2D imaging of the PFC; *U* = 37030, *p* < 0.0001, n = 515 cells for 3D imaging of the CeA, n = 231 cells for 2D imaging of the CeA; Mann–Whitney *U* test). We then measured the overlap of the spatial footprint. The spatial footprint was defined as the 50th percentile of the peak value of each spatial filter extracted by the algorithms. The overlap rate for each cell was defined as the proportion of shared pixels with other cells in its spatial footprint. We compared the spatial footprint sizes. In the recording from the PFC and CeA, the sizes of the spatial footprints were larger in 3D imaging than those in 2D (Figure 6F,G; *U* = 208000, *p* < 0.0001, n = 2083 cells for 3D imaging of the PFC, n = 1227 cells for 2D imaging of the PFC; *U* = 7117, *p* < 0.0001, n = 515 cells for 3D imaging of the CeA, n = 231 cells for 2D imaging of the CeA; Mann–Whitney *U* test). However, the overlap rate was significantly smaller in 3D imaging than that in 2D (Figure 6H,I; *U* = 347100, *p* < 0.0001, n = 2083 cells for 3D imaging of the PFC, n = 1227 cells for 2D imaging of the PFC; *U* = 29570, *p* < 0.0001, n = 515 cells for 3D imaging of the CeA, n = 231 cells for 2D imaging of the CeA; Mann–Whitney *U* test). The longer cell-cell distance, larger spatial footprint size, and smaller overlap in 3D imaging might contribute to the accurate estimation of neural activity.

## Discussion

In this study, we developed a simple, low cost 3D fluorescence microscope to use in freely moving small animals. The microscope was made from off-the-shelf optical components, CMOS image sensor electronics designed by an open-source hardware project (UCLA miniscope), and a custom 3D-printed microscope body. These parts were easily available at a low cost. Moreover, the 3D-printed parts were obtained using a consumer-grade 3D printer (Form2; Formlabs). For these reasons, the design was easy to test and modify. The weight of the microscope might affect the animal behavior and must be kept as low as possible. The microscope designed for this study weighed 3.7 g, which was acceptable for its use in mice. A further weight reduction is required to record images from smaller animals such as zebra finches ^19^. Using the new, lightweight version of the UCLA miniscope (miniscope v4; miniscope.org/index.php/Miniscope_v4) in place of the miniscope v3 might provide a suitable solution to reduce the weight. Moreover, the miniscope v4 includes an EWTL in its optical path and can capture 3D images by adding electronics for fast focus-scanning.

Not only the methods for image acquisition but also the neural activity extraction algorithms affect the quality and quantity of neuronal activity traces. However, existing algorithms for 3D imaging have not shown satisfactory performances for wide-field 3D imaging data (Figure 3B–D). Based on the CNMF algorithm ^13^, we developed a novel algorithm, CNMFw3, to extract individual cell activity from 3D images (Figure 2). The algorithm showed superior performances for simulated and real wide-field 3D imaging data (Figure 4,5). CNMFw3 uses a single, global background model and spatial high-pass filtering for rejecting background fluorescence (Figure 2B, C). CNMFw3 was significantly more performant for the simulated data than the existing methods (Figure 3B–D). However, CNMF-E uses a multiple local background model to achieve higher accuracy for wide-field 2D imaging data compared to other methods using the single background model ^18^. Therefore, adapting the local background model employed by the CNMF-E algorithm to use wide-field 3D imaging data will be an important research field for the future.

Using the new hardware and software, we could image 1.7–2.6 times more cells than with standard 2D imaging techniques (Figure 6C). Moreover, neurons extracted from 3D images showed longer cell-cell distances (Figure 6D, E), larger spatial footprints (Figure 6F,G), and smaller overlap of the spatial footprints (Figure 6H,I) than those from 2D images. Because the overlap of spatial cell footprints is a large obstacle to the neural activity extraction from wide-field imaging data ^13,18^, these features lead to a higher quality of the neural activity extraction.

Overall, by combining the newly designed hardware and software for 3D imaging, greater quality and amount of neuronal activity was recorded from freely behaving mice than with conventional methods.

## Methods

### Head-mounted 3D microscope

The miniature microscope design was based on the UCLA miniscope v3 (miniscope.org). Excitation light emitted from a blue LED (LXML-PB01-0030; Lumileds, Aachen, Germany). The light passed through an excitation filter (ET470/40x; Chroma Technology, Bellows Falls, VT) and was reflected by a dichroic mirror (T495lpxr; Chroma Technology) onto the tissue through an EWTL (Varioptic A-25H0; Corning, Corning, NY), an objective lens (aspherical singlet; #83-605, Edmund Optics, Burlington, NJ, achromatic doublet; #49-270, Edmund optics, or tandem configuration of achromatic doublets; #84-126 and #49-271, Edmund Optics), and a GRIN lens (0.5-mm diameter, 7.9-mm length; CLHS050GFT009; Gofoton, Tsukuba, Japan, or 1.0-mm diameter, 3.7-mm length CLH-100-WD002-SSI-GF3; Gofoton). The fluorescent emissions collected by the lenses were passed through the dichroic mirror and an emission filter (ET525/50m; Chroma Technology). The fluorescence image was focused by a tube lens (#49-277; Edmund Optics) and captured by a CMOS image sensor (MT9V032; ON Semiconductor, Phoenix, AZ). The microscope body was made using a 3D printer (From2, resin type FGPBLK03; Formlabs, Somerville, MA). The design files are available at https://github.com/yuichirohayashi/Zscope. The driving voltage for the EWTL (square wave, 2 kHz) was generated using a data acquisition board (PCIe-6259, National Instruments, Dallas, TX) controlled by custom software written in LABVIEW 7.1 (National Instruments) and amplified with a linear amplifier (As-904-150B, NF Corporation, Yokohama, Japan). The EWTL driver was synchronized to the image acquisition of the CMOS image sensor (Supplementaly Figure 1).

### Point spread function measurements

Three-dimensional stacks of 1-μm fluorescent beads were captured with a spacing of 10 μm. The point speed function was estimated as a Gaussian function (Figure 1E).

### Animals

All experiments complied with the relevant guidelines and regulations. Animal care and use followed protocols approved by the Animal Research Committee of Kansai Medical University. All experiments complied with the ARRIVE guidelines (https://arriveguidelines.org/arrive-guidelines/experimental-animals). Ten adult male C57BL/6N mice were used. The mice were purchased from Japan SLC, Inc. (Shizuoka, Japan), housed under a standard 12-h light/dark cycle, and allowed *ad libitum* access to food and water. At the time of the viral injection surgery, the mice were more than 10 weeks old and had body weights of 25.3 ± 0.4 g (mean ± SEM).

### Surgery

The animals were anesthetized using isoflurane. The skull was exposed, and small craniotomies (<0.5 mm) were made over the target brain regions.

AAV10-syn-GCaMP6f-WPRE (University of Pennsylvania vector core) was diluted to 5 × 10^12^ particles/mL in phosphate-buffered saline, and 200 nL were injected. The stereotaxic coordinates of the virus injection were as follows: hippocampus, AP +2.3 mm, ML 1.5 mm, and DV − 1.2 mm; PFC, AP +2.0 mm, ML 0.5 mm, and DV − 1.0 mm; CeA, AP − 1.1 mm, ML 2.5 mm, and DV − 4.5 mm. One week after the viral injection, the second surgical procedure to implant the optics proceeded. A cranial window assembly composed of a glass coverslip (0.12 mm thick) and a stainless-steel cannula (2.76-mm outer diameter, 2.40-mm inner diameter, and 1.5-mm height) for hippocampus imaging was placed over the dorsal CA1 ^20^. For PPC and CeA imaging, we implanted the GRIN lens to target these regions and glued it to the skull with dental cement (Shofu, Kyoto, Japan).

Four weeks after implantation of the optical parts, GCaMP-expressing neurons were imaged using the microscope with a tandem achromatic lens (for CA1), achromatic doublet lens (for the PFC), or aspherical singlet lens (for the CeA and PFC). The aluminum baseplate of the microscope was cemented in a position at which neurons could were visible.

### Imaging sessions

The excitation light intensity was approximately 0.1–0.5 mW/mm^2^. Images were captured at 60 Hz for 5 min. Image data were first stored in the computer’s memory before being transferred to the hard drive once the image acquisition had been completed to prevent missing frames.

### Image processing

ImageJ 1.52 (National Institutes of Health, MA) and MATLAB 2019a (Mathworks, Natick, MA) were used for all analyses. The lateral displacements of the brain were corrected by image translation using TurboReg ^21^. The registered image stream was processed with CNMFw3 or other neural activity extraction algorithms. The extracted spatial filters were then visually inspected, and those showing non-neuronal morphology were removed.

### Simulation experiments

The synthetic data used for Figure 3 were generated as follows: 100 neurons (ellipsoid of length: 4, width: 4, height: 2 pixels) were randomly placed in a volume of 100 × 100 ×10 pixels, maintaining a minimum center distance of 4 pixels in the XY axis and 1 pixel in the Z-axis. Calcium transients were generated with a Poisson process (mean transient rate 0.1 Hz, 1000-time steps at a 5 Hz frame rate) and convolved with an exponentially decaying kernel (*τ* = 0.5 s). Background fluctuations were extracted from real imaging data by moving the average filter with a window size of 10 pixels. The background was linearly mixed with the neural activity image. The volumetric data were convolved with a Gaussian kernel (SD = 2 pixels) to approximate the PSF of the microscope and scattering by the tissue. Poisson noise with a peak value of 200 was added to each pixel to simulate the photon shot noise.

## Code availability

The MATLAB implementation of CNMFw3 is available at https://github.com/yuichirohayashi/CNMFw3.

## Acknowledgements

This work is supported by JSPS KAKENHI (Grant No. 17K19436, 20K07715, 20H03547) to Y.H., JSPS KAKENHI 20H04849 to R.K.

## Author Contributions

Conceptualization, Y.H.; Methodology, Y.H.; Investigation, Y.H.; Writing –Original Draft, Y.H.; Writing –Review & Editing, Y.H., K.K. and R.K.; Funding Acquisition, Y.H. and R.K.; Resources, Y.H., K.K. and R.K.; Supervision, Y.H.

