## Supplementary Figure 1 for "Large-scale calcium imaging with a head-mounted axial scanning 3D fluorescence microscope"

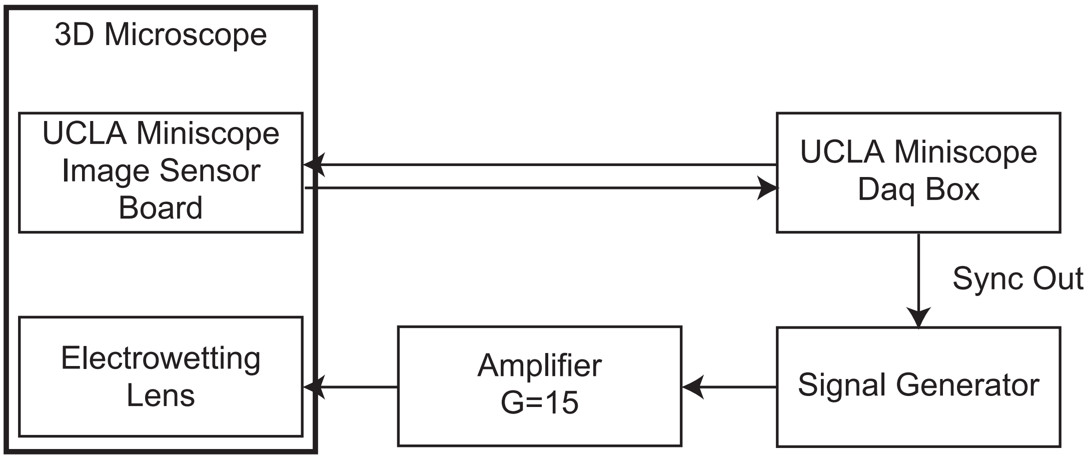


**Supplementary Figure 1. Wiring diagram for the 3D microscope system.**

The driving signal of the EWTL was triggered by the synchronized signal from the UCLA miniscope data acquisition box.
