## Supplementary Table 1 for "Large-scale calcium imaging with a head-mounted axial scanning 3D fluorescence microscope"

| | Focal Length | Numerical Aperture | Working Distance | Lateral PSF FWHM (μm) | Axial PSF FWHM (μm) |
|---|---|---|---|---|---|
| Aspherical (Asph) | 2.72 | 0.42 | 2.23 | 1.7 | 21.9 |
| Asph + 1.0 mm GRIN | - | - | - | 4.0 | 33.1 |
| Asph + 0.5 mm GRIN | - | - | - | 5.8 | 76.1 |
| Achromatic Doublet (Ach) | 4.5 | 0.28 | 2.99 | 5.7 | 18.4 |
| Tandem Achromatic Doublets (TAch) | 3.55 | 0.35 | 1.42 | 1.9 | 15.9 |

**Supplementary Table 1. Optical parameters of the objective lenses**

Focal length, numerical aperture, and working distance were calculated using the OpTaliX version 8.70 software.

FWHM = full-width at half-maximum; PSF = point spread function
